# Glioblastoma gene expression based subtypes have defined metabolomic states

**DOI:** 10.1101/2025.09.05.674480

**Authors:** Hélèna L. Denis, Jaëlle Méroné, Akhil Shukla, Valérie Watters, Victoire Fort, Gabriel Khelifi, Mikalie Lavoie, Line Berthiaume, Félix Rondeau, Émy Beaumont, Karine Michaud, Myreille D’Astous, Stephan Saikali, Marc-Étienne Huot, Étienne Audet-Walsh, Maxime Richer, Samer M.I. Hussein

**Affiliations:** Université Laval, Québec, QC, Canada; Axe neurosciences du Centre de recherche du Centre hospitalier universitaire (CHU) de Québec-Université Laval, Québec, QC, Canada; Axe oncologie du Centre de recherche du CHU de Québec-Université Laval, Québec, QC, Canada; Axe endocrinologie et néphrologie du Centre de recherche du CHU de Québec-Université Laval, Québec, QC, Canada; Department of Surgery, CHU de Québec-Université Laval, Québec, QC, Canada; Department of Molecular Biology, Biochemistry and Pathology, CHU de Québec, Université Laval, Québec, QC, Canada; Department of Molecular Medicine, CHU de Québec, Université Laval, Québec, QC, Canada

**Keywords:** human glioblastoma stem cells, biobank, transcriptome, metabolomics, gene expression, isocitrate dehydrogenase

## Abstract

Glioblastoma (GBM) is a highly aggressive primary brain cancer with poor prognosis (<15 months), highlighting the urgent need for more effective therapies. As current treatments are not effective, the need for a deeper understanding of the biology of GBM cells, including how they reprogram their metabolism to support their aberrant and uncontrolled growth, is critical. To this end, we established a collection of 41 human glioma cell lines derived from freshly resected tumour tissues from 99 patients. We characterized 12 of these cell lines by combining histologic, genetic, stem cell derivation and self-renewal, and metabolomic analyses. Histological and genetic profiles included IDH mutation status, Ki-67 proliferation index, ATRX status, mutant *TP53* expression, chromosome 10q loss, *EGFR* amplification, and MGMT promoter methylation. Of these, only p53 mutation expression status showed weak segregation of the cell lines into 2 separate metabolic groups based on amino acid levels, but none showed an effect on stem cell derivation or self-renewal. Further characterization of these 12 cell lines revealed significant metabolic and phenotypic differences when comparing mesenchymal versus proneural gene expression subtyping. We show significant increases in TCA cycle metabolites in mesenchymal-like GBM cells and higher overall metabolic activity compared to proneural-like cells. These findings highlight the complexity of GBM and the need for personalized treatments that consider the metabolome of each subtype as a potential therapeutic avenue.

## INTRODUCTION

Gliomas, glioneuronal tumours, and neuronal tumours represent the most frequent and diverse tumours affecting the central nervous system (CNS) [1]. According to the fifth edition of the World Health Organization (WHO) classification of CNS tumours [2], they are divided into six distinct groups: (1) Adult-type diffuse gliomas include glioblastoma (IDH-wildtype), astrocytoma (IDH-mutant), and oligodendroglioma (IDH-mutant and 1p/19q-codeleted), which are commonly encountered in adult neuro-oncology practice; (2) Pediatric-type diffuse low-grade gliomas are typically associated with a favorable prognosis; (3) Pediatric-type diffuse high-grade gliomas are generally more aggressive; (4) Circumscribed astrocytic gliomas are characterized by a more localized growth pattern compared to the infiltrating nature of adult-type diffuse and pediatric-type diffuse high-grade gliomas; (5) Glioneuronal and neuronal tumours, a diverse group often exhibiting neuronal differentiation, along with (6) ependymal tumours, represent the remaining categories in the classification.

Among adult-type diffuse gliomas, IDH-wildtype (wt) glioblastoma (GBM) are the most common and aggressive primary brain tumours in adults [1]. Their highly infiltrative and recurrent nature leads to failures following chemotherapy and radiotherapy, as well as a short median survival (less than 15 months) [3]. There is an urgent need to develop new therapies to improve the survival and quality of life of individuals affected by GBM. IDH-wt GBM exhibits molecular heterogeneity, evident not only between different tumours (intertumoral heterogeneity) but also within the same tumour (intratumoral heterogeneity). This variability reflects the diversity in its histological and molecular characteristics, cellular origins, topographical variations, growth patterns, and therapy sensitivities, all contributing to its aggressive behavior and poor prognosis. Key molecular alterations in GBM include *TERT* promoter mutations, *EGFR* gene amplification, and the combined gain of chromosome 7 and loss of chromosome 10 (+7/−10) [2]. GBMs typically lack ATRX mutations, while TP53 mutation status and MGMT promoter methylation are key factors influencing prognosis and treatment response[2]. Comprehensive transcriptomic analyses further delineated this heterogeneity, identifying molecular signatures that categorize various adult-type GBM subtypes, with one subtype in particular, the mesenchymal-like subtype, which is associated with increased invasive properties, poor prognosis, and low patient survival rate [4–8].

Additionally, the metabolism of GBM cells also shows significant heterogeneity and plasticity. While these cells primarily use glucose as their main metabolic fuel, they can also utilize other substrates depending on their genetic makeup and environmental conditions. These include amino acids such as glutamine and glutamate, lipids like fatty acids and lipid droplets, and other carbon sources such as acetate [9]. The robust proliferative activity of cancer cells demands a high supply of energy in the form of ATP, precursors for macromolecules such as carbohydrates, lipids, proteins, and nucleic acids, and reducing equivalents to support their rapid growth and survival. Furthermore, these cells continuously adapt their metabolism to their ever-changing environmental conditions [10].

Given the limited success of traditional cancer therapies targeting mechanisms such as growth, cell cycle regulation, autophagy, DNA repair, apoptosis, angiogenesis, and immune checkpoints, it is increasingly crucial to advance our understanding of oncogenic metabolic mechanisms and their associated signaling pathways to identify innovative treatment strategies [11–13]. While high-throughput data from GBM tissues and cell lines have been extensively shared, there are only a few recent databases that include both tissues and cell lines [14–17]. In alignment with this goal, we established a collection of human glioma cell cultures derived from freshly resected tumour tissues from patients undergoing surgery. We cultured these cells under conditions that preserve glioma stem cells (GSCs). This is particularly important as GSCs are thought to drive recurrence and therapy resistance due to their unique capabilities for growth, progression, and resilience against treatments [18–20]. We then performed metabolomic analysis on 12 IDH-wt GSC lines, with different histopathological and genetic properties, which include mitotic activity, *TP53* mutation expression, *MGMT* promoter methylation, 10q loss and *EGFR* amplification. We found no clear correlation in their metabolomes and clinical pathology diagnosis, except for *TP53* mutation where we observe a marked increase in amino acid levels when *TP53* is mutated. Moreover, the frequency of GSC derivation and their self-renewal was also not affected by clinical pathology diagnosis. However, when GSCs are segregated by gene expression-based GBM subtyping, we observe a significant increase in TCA intermediates specifically in mesenchymal-like GSC subtypes, the most aggressive GBM subtype. Our results suggest that metabolomic changes align with gene expression-based subtyping and may lead to better patient diagnosis compared to just histopathological and genetic testing.

## RESULTS

### Human glioma biobank establishment and cell culture collection

To establish a continuous, local, and reliable glioma patient tissue samples, glioma surgical resection specimens were collected, and each sample was divided into three distinct pieces (**Fig. 1a**). The first piece was flash-frozen to preserve the molecular integrity of the tissue. The second piece was processed into formalin-fixed, paraffin-embedded (FFPE) blocks for histopathological and molecular analyses. The third piece underwent mechanical and enzymatic dissociation to isolate single cells. A total of 99 patients were included in the cohort, 18 IDH-mutant gliomas (13 astrocytomas and 5 oligodendrogliomas) and 81 IDH-wildtype glioblastomas (**Fig. 1a**). The group comprised 43 women and 56 men, with a mean age of 59 years (**Fig. 1b**). The majority of cases (76%) were derived from initial resections performed at the time of primary diagnosis, prior to the administration of any treatment (**Fig. 1c**). The cohort included a higher number of patients with IDH-wt status (82%), which is consistent with what is observed in the U.S.A. [21]. As expected [2], the Ki-67 proliferative index was higher in IDH-wt GBM (**Fig. 1d**), with *ATRX* status aligning accordingly (86% vs 14%; **Fig. 1e**). Additionally, 57% of tumours exhibited *TP53* overexpression, suggestive of a *TP53* gene mutation (**Fig. 1e**). Notably, *TP53* overexpression was identified in 35% of IDH-wt cases compared to 50% of IDH-mutated cases. *MGMT* promoter hypermethylation was observed in 53% of IDH-wt samples (**Fig. 1e**), and IDH-mutated biopsies were not tested for *MGMT* status. Thus, we established a large biobank of glioma samples and cell lines that can be exploited to further our knowledge on glioma biology.

**Figure 1.**
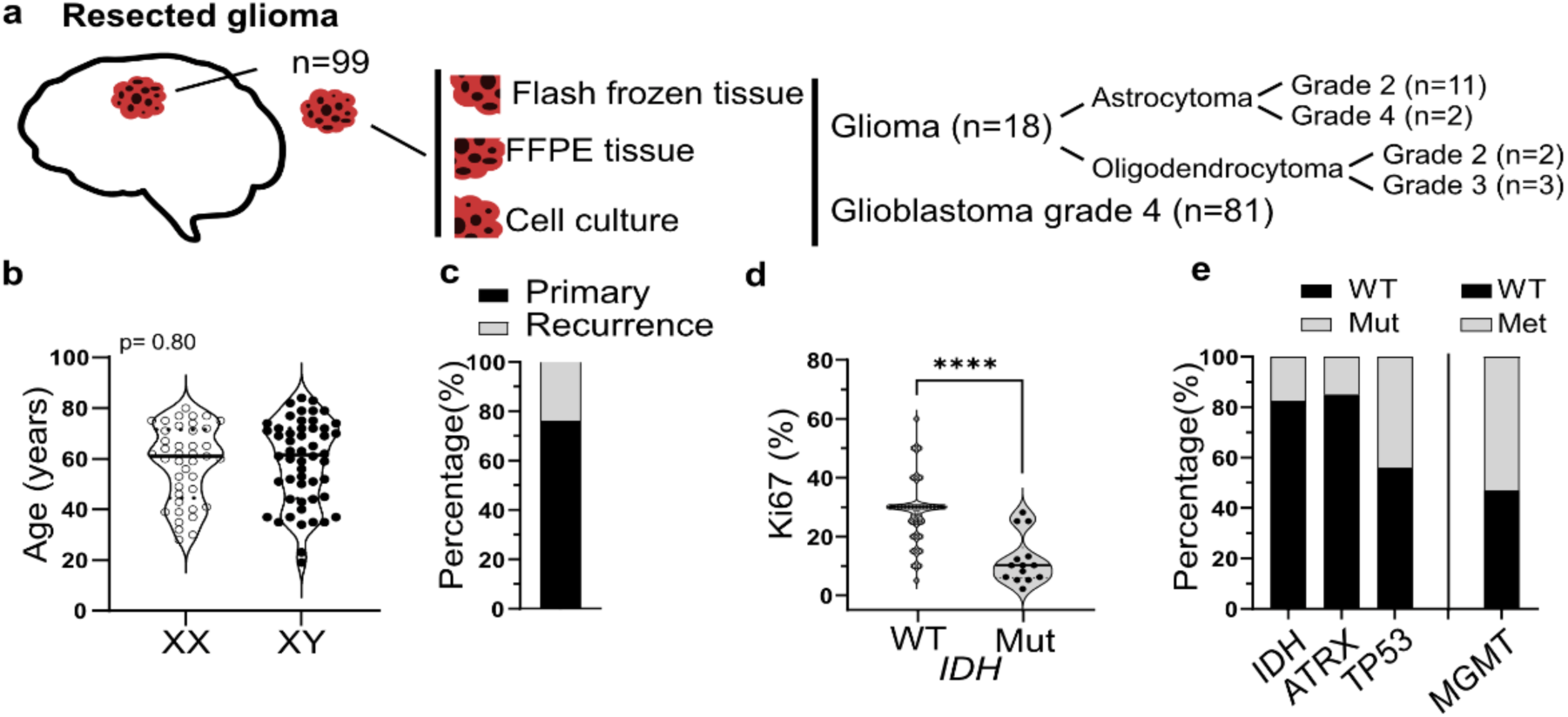
Human glioma biobank establishment and cell culture collection. **a.** Generation of the glioma biobank from resected human tissue. **b.** Biopsies came from 43 women and 56 men with a similar age (mean= 59 years old). **c.** Most samples originated from a primary resection. **d.** Percentage of Ki67-positive cells within the biopsy compared to IDH status. **e.** Percentage within our biobank of biopsies with WT or Mut status. Most of the biopsies are *IDH* (82%), *ATRX* (86%) and *TP53* (57%) WT. Half of the biopsy are *MGMT* WT. Mutated IDH biopsies are usually not tested for *MGMT* status. *<u>Statistical analysis:</u>* **d.** unpaired t-test. *<u>Abbreviations:</u> ATRX*, X-linked nuclear protein; FFPE, Formalin-Fixed Paraffin-Embedded; *IDH*, isocitrate dehydrogenase; n, number; *MGMT*, O^6^-methylguanine-DNA methyltransferase; Met, methylation; Mut, mutated; *TP53*, tumor protein 53; XY, men; XX, women; WT, wild-type.

### Establishment and Characterization of Patient-Derived GBM Cell Lines

The initial cohort was representative in terms of age, gender, and the distribution of molecular alterations, with no evidence of selection bias. From this cohort, part of the biopsy underwent dissociation processing (**Fig. 2a**) and was cultured to obtain and calculate success in GSCs sphere formation (**Fig. 2b**). Among the glioma IDH-mutant samples, 3 did not form spheres, 4 formed spheres within one month, and 11 after more than one month (**Fig. 2b**). Monolayer cultures were successfully established from only three long-term sphere-forming glioma samples. For GBM samples, 39 did not form spheres, 18 formed spheres within one month, and 24 after more than one month. Monolayer cultures were established for 38 samples (**Fig. 2b**). Among the 38 GBM cases that successfully generated cell lines, there were 17 women and 21 men, with mean ages of 66 and 62 years, respectively (**Fig. 2c**). Of these, 86% of the cases originated from primary resections, while the rest were recurrent tumours (**Fig. 2d**). GBM cell lines had an average Ki67 index of 30% and *ATRX*-wt status as expected [2] (**Fig. 2e**). The distribution of *TP53* status was 64% WT and 36% Mut, while *MGMT* promoter methylation was 48% WT vs 52% Met (**Fig. 2f**). These 38 cell lines thus strongly represent the distribution of histopathological and genetic profiles found in our tissue biobank for GBM samples.

**Figure 2.**
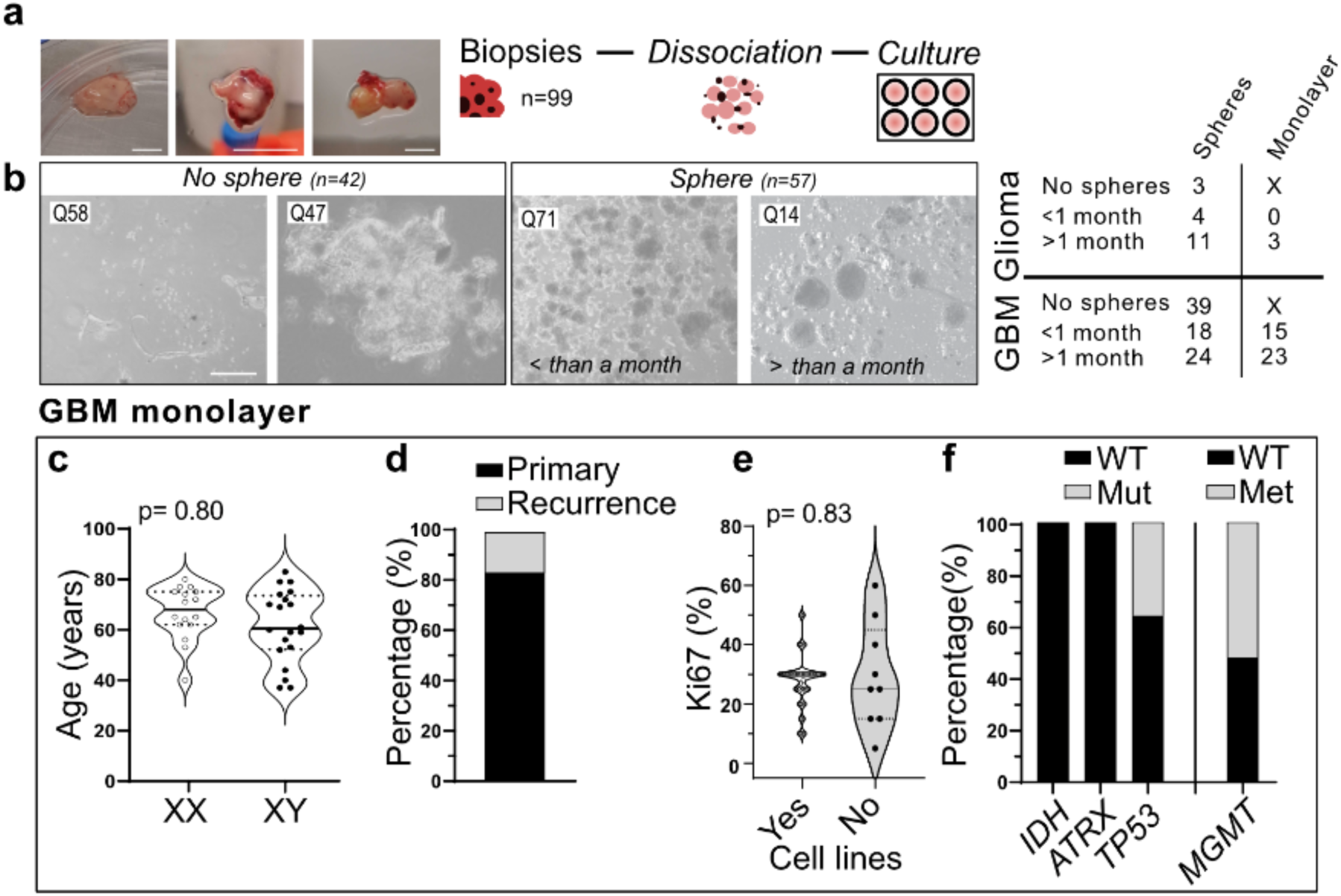
Establishment and characterization of patient-derived GBM cell lines. **a.** Images of biopsies before dissociation for culture. **b.** Proportion of glioma and GBM biopsies that generated spheres or monolayers versus those that did not. **c.** GBM cell lines came from 17 women and 21 men with a similar age (women, 66 years old; men, 62 years old). **d.** Cell lines mostly came from a primary resection. **e.** Approximately 30% of cells in the corresponding biopsies are Ki67-positive, with no difference observed between biopsies that generated cell lines and those that did not. **f.** All our cell lines are *ATRX* WT. Percentage within our cell lines with WT or Mut/Met status: 53% and 45% of the cell lines are *TP53* and *MGMT* WT respectively. *<u>Scale bar:</u>* **a.**1 cm **b.** 200µm. *<u>Statistical</u> <u>analysis:</u>* Unpaired t-test or Mann-Whitney test. *<u>Abbreviations:</u> ATRX*, X-linked nuclear protein; *EGFR*, epidermal growth factor receptor; GBM, glioblastoma; *IDH*, isocitrate dehydrogenase; Loss chr10q, loss of the chromosome 10q; *MGMT*, O^6^-methylguanine-DNA methyltransferase; Mut, mutated; *TP53*, tumor protein 53; XY, men; XX, women; WT, wild-type.

### GBM cell line establishment is not affected by clinical pathology

It is thought that GSCs can recapitulate GBM biology and help study and understand the progression of GBM. To address this, we further characterized 12 individual GBM GSC lines as they relate to their clinical pathology (**Fig. 3**). We observed morphological differences among these lines, with most showing a fibrillary morphology in the adherent state, as expected from astrocytic-like cells (**Fig. 3a**). A limiting dilution assay, used to evaluate the self-renewal and proliferative capacity of GSCs, revealed significant variability, with GSC percentages ranging from 4.5% to 26% (**Fig. 3b**). Moreover, among these 12 cell lines, no correlations were observed between GSC percentage and factors such as sex (although the number of samples was limited), age, *TP53* status, *MGMT* status, *EGFR* amplification, or loss of chromosome 10q (**Fig. 3c**). These results demonstrate factors that are not linked to patient clinical pathology may affect GSC derivation and maintenance. Additionally, and since GSCs are postulated to promote GBM disease progression, other characteristics and profiles are needed to fully diagnose GBM.

**Figure 3.**
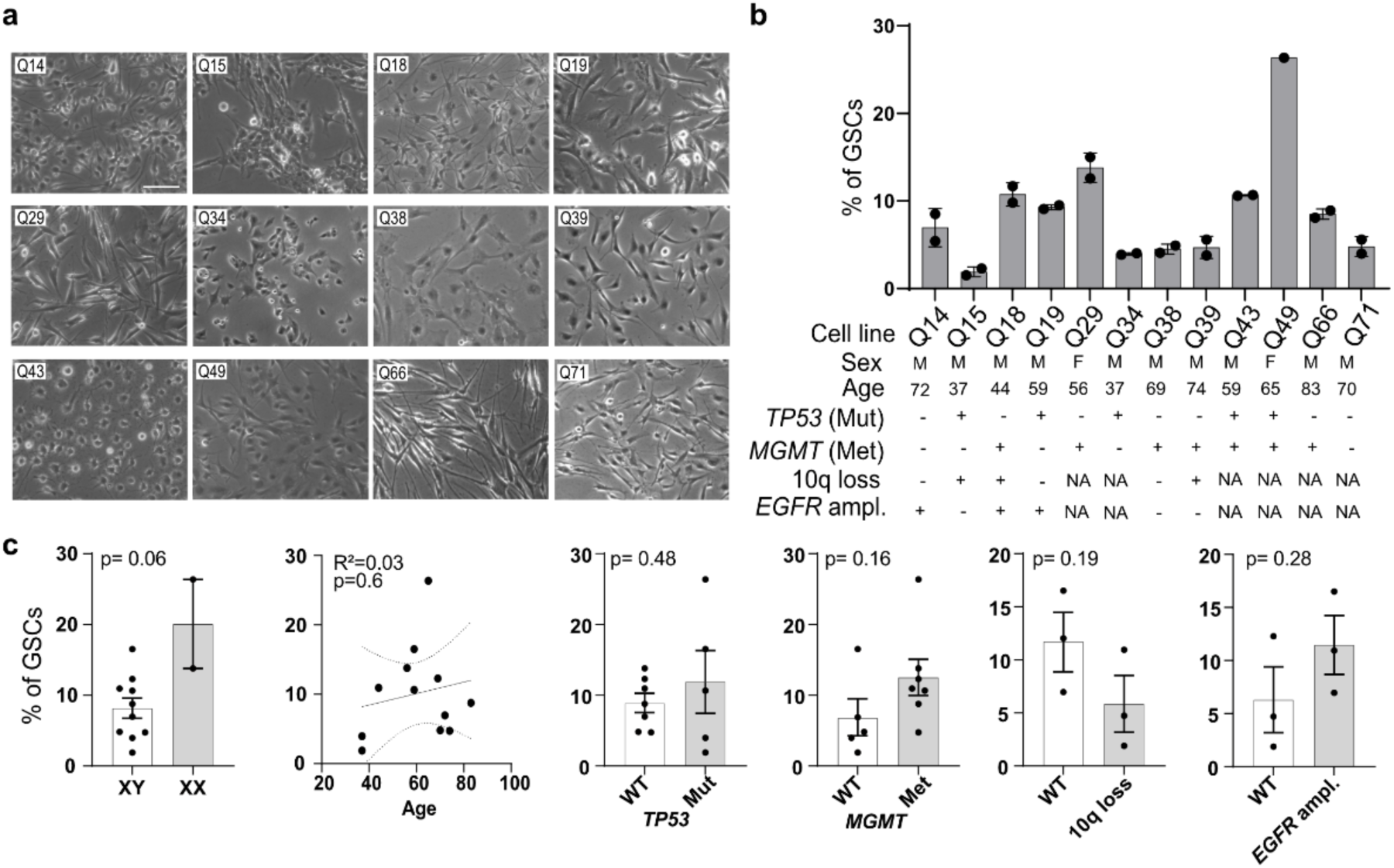
GBM cell line establishment is not affected by clinical pathology. **a.** Pictures of 12 GSC cell lines to be used in the downstream experiments. **b.** Using limiting dilution assay, we assessed the percentage of GSCs within the 12 cell lines. **c.** No correlations have been found between GSC percentage and sex, age, *TP53*, *MGMT*, loss of the chromosome 10q status, *EGFR* amplification. *<u>Scale bar:</u>* 100µm. *<u>Statistical analysis:</u>* Unpaired t-test or Mann-Whitney test; simple linear regression followed by Pearson correlation test. *<u>Abbreviations:</u>* ampl., amplification; *EGFR*, epidermal growth factor receptor; GSCs, glioblastoma stem cells; Met, methylated; *MGMT*, O^6^-methylguanine-DNA methyltransferase; Mut, mutated; *TP53*, tumor protein 53; WT, wild-type; XY, men; XX, women; 10q loss, loss of the chromosome 10q.

### *TP53* mutations alter metabolomic profiles in GSC lines

As mentioned above, histopathological and genetic testing alone cannot distinguish between GSC derivation potential and characteristics, which may reflect the difficulty in segregating individual GBM differences using current diagnostic strategies. To answer this need, we performed metabolomic profiling on the 12 GBM GSC lines as an additional method to integrate with clinical pathology. We found lines harboring *TP53* mutations showed higher amino acid levels compared to *TP53* wt lines (**Fig. 4c**). Principal component analysis (PCA) identified modest segregation at sample level (**Fig. 4a**), but revealed distinct metabolomic signatures associated with *TP53* mutation status at the level of individual metabolites. PCA score plots showed partial clustering of *TP53* mutant and wild-type cell lines, with 95% confidence ellipses indicating significant overlap **(Fig. 4a).** This suggests that no single combination of metabolites can fully differentiate these groups. However, PCA biplots highlighted specific metabolite families, including leucine, isoleucine, valine, glutamate, aspartate that were consistently associated with *TP53* mutations **(Fig. 4b).** Further analyses of TCA cycle intermediates and other metabolites did not reveal clear differences according to *TP53* status (**Suppl. Fig. S1a-b**). In contrast, no differences in metabolite levels were observed based on *MGMT* methylation status, as both PCA score plot and biplot analyses failed to reveal clear separation between *MGMT* wt and methylated cell lines (**Fig. 4d-e**, **Suppl. Fig. S1c-e**). Metabolite profiling was also performed according to chromosomal and genetic alterations. Cell lines with 10q loss (**Suppl. Fig. S2**) or *EGFR* amplification (**Suppl. Fig. S3**) did not show major changes in amino acid levels, TCA intermediates, or other metabolites compared with lines without these alterations, suggesting that *TP53* status has a stronger impact on metabolic profiles in GBM cell lines than MGMT methylation, 10q loss, or EGFR amplification.

**Figure 4.**
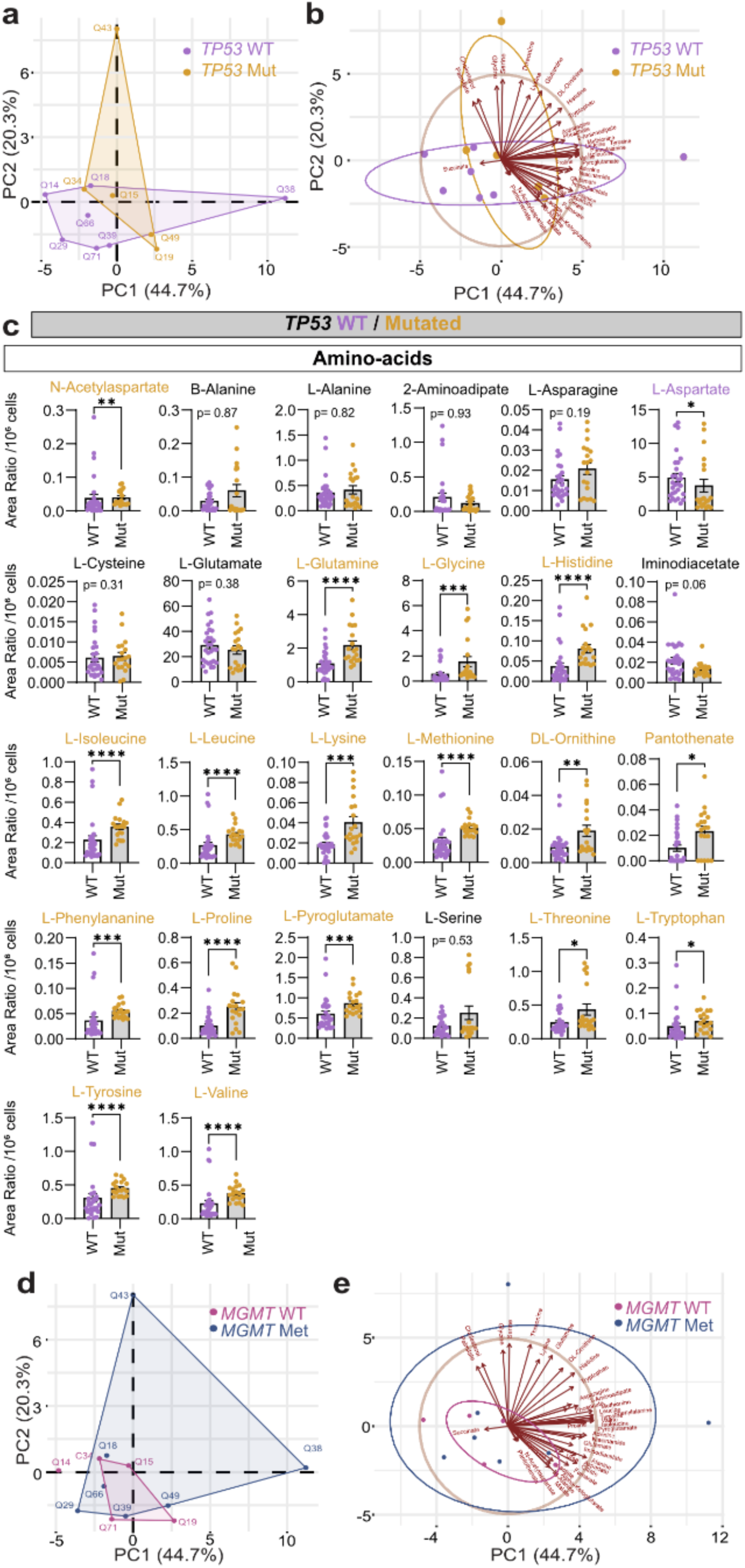
*TP53* mutations alters metabolomic profiles in GSC lines. Metabolomic profiling revealed higher amino acid levels in glioblastoma cell lines with *TP53* mutations compared to wild type (**a-e**), whereas no differences were observed in relation to *MGMT* mutation status (**f-g**). **a.** PCA score plot did not show a clear separation of the cell lines according to *TP53* status. **b.** PCA biplot showed that some metabolites levels are associated with *TP53* status. **c.** Levels of amino acids are increased in cell lines with *TP53* mutations. **d,e.** PCA score and biplot analyses did not reveal a clear separation of cell lines based on *MGMT* methylation status. *<u>Statistical analyses:</u>* Confidence ellipse (95%) highlights group clustering, and the arrows indicate the contribution and direction of individual metabolites to the observed separation between the subtypes. Unpaired t-test or Mann-Whitney test. *<u>Abbreviations:</u>*AA, amino-acids; Met, methylated; *MGMT*, O^6^-methylguanine-DNA methyltransferase; Mut, mutated; *TP53*, tumor protein 53; WT, wild-type.

### GBM subtypes show distinct metabolomic states

Since clinical pathology diagnoses did not clearly distinguish metabolic profiles, we opted to use gene expression-based classification to assess whether previously identified GBM molecular subtypes [8] could be associated with distinct metabolic patterns in GBM GSC lines. To classify our cell lines, RNA expression of the 12 selected GSCs was analyzed by RT-qPCR (**Fig. 5**). Six genes were selected as markers of the mesenchymal-like subtype (*BMI1*, *CD44*, *SNAI2*, *TNFSF10*, *ANXA1*, and *ANXA2*), and four genes as markers of the proneural-like subtype (*OLIG2*, *PDGFRA*, *ASCL1*, and *ERBB3*). Hierarchical clustering of the expression profiles segregated the lines into two groups, with eight classified as mesenchymal-like and four as proneural-like (**Fig. 5a**). No correlations were found between proneural-like and mesenchymal-like cell lines in terms of GSC percentage, age, Ki67 percentage and clinical features, including *TP53*, *MGMT*, loss of the chromosome 10q status and *EGFR* amplification (**Fig. 5b-e**), again highlighting that clinical pathology alone is not sufficient to classify GBM diagnosis. However, PCA plots of the metabolomic profiles revealed distinct clusters for proneural-like (blue) and mesenchymal-like (red) samples, indicating that the metabolome of these GBM cells is distinct between subtypes (**Fig. 6a**). Indeed, the metabolic profile of these GBM GSC lines allowed complete separation of proneural-like from mesenchymal lines. Next, we performed correlation analyses to better understand the metabolic profile of the two studied GBM GSC line phenotypes. In proneural-like lines, a strong correlation cluster can be observed for amino acids, which is mostly non-correlated or inversely correlated with TCA cycle intermediates and other metabolites (**Fig. 6b**, left). In mesenchymal lines, while the high correlation between amino acids is also observed, these molecules also positively correlated with the other studied metabolites, such as TCA cycle intermediates (**Fig. 6b**, right), suggesting a distinct mitochondrial metabolic profile in these cancer cells. Indeed, mesenchymal-like lines were characterized by a stronger signature of TCA cycle intermediates such as citrate, fumarate, and malate (**Fig. 6c**). Moreover, these analyses showed a high abundance in proneural-like lines of one-carbon metabolism amino acids (serine, glycine) and lipid metabolites (cholesterol, palmitate) (**Suppl. Fig. S4**). These results indicate that proneural-like and mesenchymal-like GBM cells have distinct metabolic programs.

**Figure 5.**
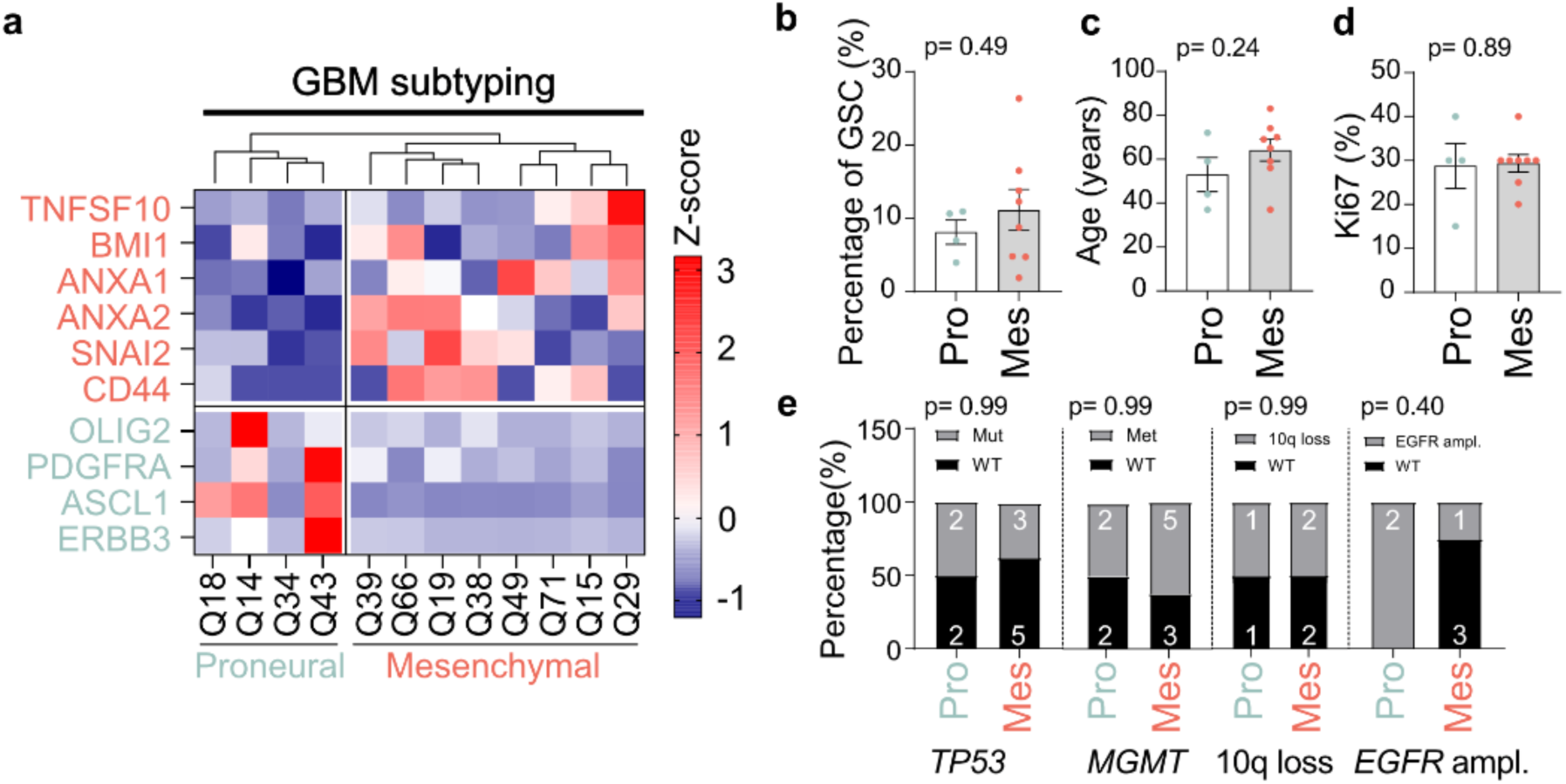
GBM cell line subtyping is not affected by clinical pathology. **a.** RT-qPCR was performed using six Mes-specific primers and four Pro-specific primers. Hierarchical clustering of the expression profiles separated the twelve cell lines into four proneural-like and eight mesenchymal-like subtypes. No correlations have been found between proneural-like and mesenchymal-like cell lines with GSC percentage (**b**) and age (**c**), Ki67 percentage (**d**) and clinical features (**e**) including *TP53*, *MGMT*, loss of the chromosome 10q status and *EGFR* amplification. *<u>Statistical analysis:</u>* **b-d**. Unpaired t-test or Mann-Whitney test; **e.** Fisher’s exact test. *<u>Abbreviations:</u>*ampl., amplification; *EGFR*, epidermal growth factor receptor; GBM, glioblastoma; GSCs, glioblastoma stem cells; Mes, mesenchymal-like cells; Met, methylation; *MGMT*, O^6^-methylguanine-DNA methyltransferase; Mut, mutated; *TP53*, tumor protein 53; Pro, proneural-like cells; WT, wild-type; 10q loss, loss of the chromosome 10q.

**Figure 6.**
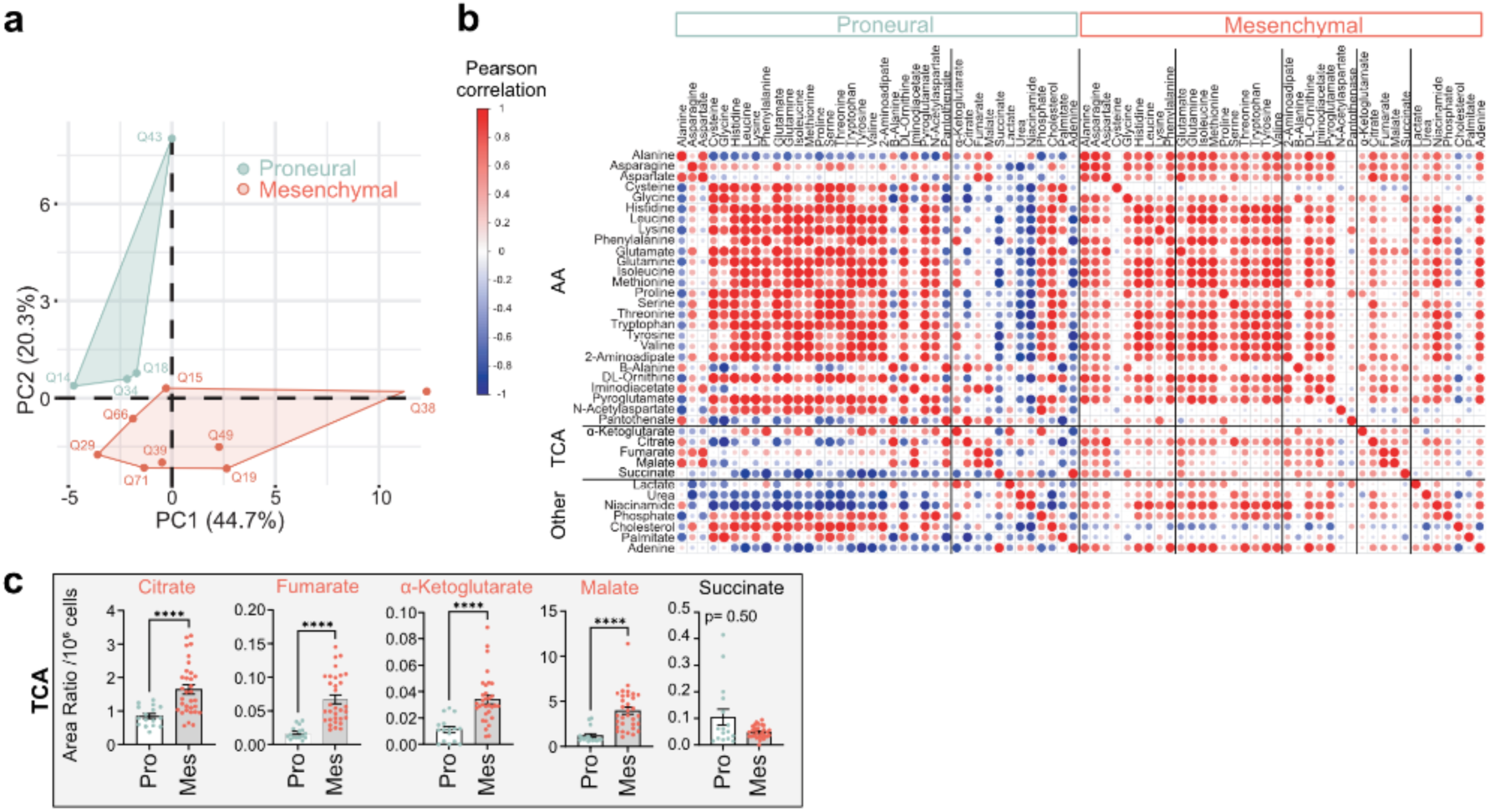
GBM subtypes show distinct metabolomic states. **a.** PCA score plot showing the separation of proneural-like (blue) and mesenchymal-like (red) cell lines. **b.** Correlation coefficient heatmaps for proneural-like (left) and mesenchymal-like (right) subtype samples showing pairwise Pearson correlation coefficients between all metabolites. **c.** Comparison between proneural-like and mesenchymal-like metabolite levels for TCA intermediates. *<u>Statistical analysis:</u>***b.** Heatmap clustering was performed using the Average Linkage method with Pearson correlation as the distance metric. The color intensity indicates the correlation strength, from −1 (blue, strong negative correlation) to +1 (red, strong positive correlation). **c.** Unpaired t-test or Mann-Whitney test. *<u>Abbreviations:</u>* AA, amino acids; Mes, mesenchymal-like cells; Pro, proneural-like cells; TCA, Tricarboxylic Acid Cycle.

## DISCUSSION

At the molecular level, GBM is highly heterogeneous. The origin of GBM tumours is still unknown, but molecular drivers have been identified. The most common mutations are found in *EGFR, TP53* and *ATRX* genes [2]. Despite this, GBM show high inter- and intratumoral heterogeneity at genetic, transcriptional, functional, and spatial levels [22–24]. Tumours may contain therapy-resistant GSCs, more sensitive differentiated cells, and genetically distinct subclones – contributing to recurrence and treatment failure [5]. GBM cells, particularly those with stem-like properties, also possess the ability to undergo metabolic reprogramming to meet the increased energy, lipid, protein, and sugar demands required for their growth, proliferation, and motility. This metabolic plasticity is essential for sustaining the self-renewal and tumorigenic potential of GSCs, highlighting the complex interplay between the stem cell-like characteristics of GBM cells and their metabolic needs for tumour progression. Recognizing the critical role of GSCs in the aggressiveness and therapy resistance of glioblastoma [18–20,25], we focused our efforts on establishing new GSC lines derived from GBM patient biopsies. GBM cells, particularly those with stem-like properties, also possess the ability to undergo metabolic reprogramming to meet the increased energy, lipid, protein, and sugar demands required for their growth, proliferation, and motility. This metabolic plasticity is essential for sustaining the self-renewal and tumorigenic potential of GSCs, highlighting the complex interplay between the stem cell-like characteristics of GBM cells and their metabolic needs for tumour progression. Our findings suggest that GSC percentage is independent of patient age, sex, or clinical pathology features, including *TP53* and *MGMT* status, as well as genetic alterations such as chromosome 10q loss and *EGFR* amplification. However, metabolomic profiling of the 12 GBM GSC lines highlighted a potential link between *TP53* mutations and elevated amino acid levels (**Fig. 4**). These results indicate that the influence of *TP53* mutations may be more pronounced on amino acid metabolism rather than central carbon metabolism. In contrast, *MGMT* methylation, 10q loss, and *EGFR* amplification did not significantly influence amino acid levels, TCA intermediates, or overall metabolite profiles. Together, these results suggest that *TP53* status may exert a more specific and detectable effect on the metabolic phenotype of GBM cells than other frequently observed genetic alterations. This finding underscores the importance of considering *TP53*-mediated metabolic reprogramming in GBM and may have implications for targeted metabolome in *TP53*-mutant GBM.

Over the past 20 years, multiple groups have conducted analyses to further characterize GBM heterogeneity based on transcriptome. These studies, including TCGA, initially identified four subtypes (classical, mesenchymal, proneural and neural) [8]. Some of these subtypes also have clinical relevance: mesenchymal GBM subtypes are linked to poor outcomes, while proneural GBM subtypes faring better [26–28]. Our classification was based on these two subtypes by targeting genes specifically expressed in each group, as identified by recent single-cell analyses [5]. Our results show that the twelve cell lines can be distinguished into these two subtypes. It is important to emphasize the significance of these subgroups. Indeed, subtype transitions can occur; for example, proneural-to-mesenchymal transition (PMT) has been observed in ∼30% of recurrences, promotes therapy resistance [27,28], and is driven by microenvironmental signals activating NF-κB [4,29–31]. These findings highlight that subtype plasticity and PMT further contribute to tumour heterogeneity and therapeutic failure [31]. Our results also show that the metabolic profiles of these two subgroups is distinct (**Fig. 6**), allowing the separation of GBM cell lines based on their subtypes. A clear characterization based on both the metabolism and transcriptome is still lacking. One study has shown that mesenchymal GBM exhibits significantly higher glycolytic activity compared to proneural GBM, which is in line with our results showing higher lactate levels in mesenchymal cells (**Fig. S4e**) [32]. A recent study combining single-cell RNA sequencing with spatial metabolomics on tissue sections (n = 6 patients) identified that a reactive-hypoxia subgroup marked by hypoxia-response genes (e.g., VEGFR, HMOX1, GAPDH) is associated with distinct glycolytic metabolic program (e.g., LDHA, PGK1) [33]. This subgroup showed the strongest overlap with the hypoxia-mesenchymal-like state defined by Neftel et al., 2019 [5]. Metabolic enrichment analyses further resolved three metabolomic subgroups within this reactive-hypoxia subgroup: subgroup 1 - pentose phosphate pathway; subgroup 2 - phosphoadenylate metabolism, a hallmark of glioma metabolism [34]; and subgroup 3 - glycolysis and amino sugar metabolism. Furthermore, the baseline transcriptional state of GBM cells corresponds to AC-, OPC-, or NPC-like [35–37] and is spatially associated with lower cellular density and enrichment of the pentose phosphate pathway [33]. Tumor growth induces nutrient and oxygen deficiency, triggering a hypoxia-driven switch from pentose phosphate pathway to glycolysis, which promotes the go/migrate program, S-phase arrest, and accumulation of de novo copy-number alterations [38,39]. By integrating metabolic characterization with the gene expression-based classification of GBM, vulnerabilities of each GBM can be pinpointed, and personalized treatment options can be explored.

Our results also show an increase in palmitic acid levels in proneural-like cell lines. This is interesting as previous studies have identified palmitic acid and oleic acid as the predominant fatty acids in glioblastoma, highlighting their central role in lipid metabolism that supports tumour growth and progression [40,41]. Palmitic acid contributes to membrane synthesis and protein palmitoylation, processes critical for cell proliferation and signaling. The strong negative correlation between citrate and palmitate in proneural cells might suggest that citrate, in addition to fueling the TCA cycle, is also exported out of the mitochondria to feed the *de novo* synthesis of fatty acids, thus increasing palmitate synthesis. Future mechanistic studies are required to fully understand the different metabolic adaptations that occur in GBM cells, and how they differ or are similar between GBM subtypes, such as between proneural-like and mesenchymal-like cells.

Taken together, our results suggest that the combination of mutations and transcriptome profile in a GBM may create unique metabolic signatures that contribute to tumour progression, resistance to therapy, and disease outcomes, while also creating specific metabolic vulnerabilities that could be exploited for therapeutic purposes. Therefore, it is essential to conduct further studies to better understand how these mutations interact and how they drive metabolic reprogramming in GBM.

## METHODS

### Ethics statement and participant recruitment

To create our human GBM cell culture resource, freshly resected tumour tissues were collected from patients undergoing surgery at the CHU de Québec – Université Laval, Hôpital de l’Enfant-Jésus. The institutional review board (#2023-6392) approved the study in accordance with the Declaration of Helsinki, and all participants provided written informed consent. Our clinical pathological facilities at CRCHU de Québec provided results for: 1) loss of 10q chromosome, assessed by PTEN loss, and amplification of EGFR using FISH, and 2) p53, IDH and ATRX presence or absence of mutation through immunohistochemistry.

### Cell culture media

#### GBM media

NeuroCult™ NS-A Basal Medium (StemCell Technologies, #05750) supplemented with 2 µg/mL Heparin (Sigma, #H3393), 10 ng/mL FGF2 (PeproTech, #100-18B), 20 ng/mL EGF (PeproTech #AF-100-15) and 1µM CHiR (Cedarlane Labs, #S2924).

#### DMEM-F12 media

DMEM/F-12, HEPES (Thermofisher Scientific Gibco™, #11330057) supplemented with 20mM of GlutaMAX™ Supplement (Thermofisher Scientific Gibco™, #35050061) 1x of MEM Non-Essential Amino Acids Solution (Thermofisher Scientific Gibco™, #11140050), 1mM of Sodium Pyruvate (Thermofisher Scientific Gibco™, #1360070), 0.11 mM of 2-Mercaptoethanol (Thermofisher Scientific Gibco™, #21985023), 40mg/mL of human recombinant Insulin (Thermofisher Scientific Gibco™, #12585014) and 1x of N2 (100x, homemade and composed of progesterone (6 mg/mL), putrescine dihydrochloride (160 mg/mL), Sodium selenite 30 mM, apo-Transferrin human (100 mg/mL).

#### Neurobasal media

Neurobasal™ Medium Catalog (Thermofisher Scientific Gibco™, #21103049) supplemented with 20mM GlutaMAX™ Supplement (Thermofisher Scientific Gibco™, #35050061), 1x of MEM Non-Essential Amino Acids Solution (Thermofisher Scientific Gibco™, #11140050) and 1x of B-27™ Supplement (Thermofisher Scientific Gibco™, #17504044).

#### Neurosphere media

2 portions of DMEM-F12 media for 1 portion of Neurobasal media supplemented with 500 ng/mL Heparin (Sigma, #H3393), 20 ng/mL FGF2 (PeproTech, #100-18B) and 20 ng/mL EGF (PeproTech #AF-100-15).

### Preparation of GBM cell lines

The biopsy was divided into three fragments (**Fig. 1a**). One fragment was flash-frozen and stored in liquid nitrogen, while another was formalin-fixed and paraffin-embedded. The last fragment was processed under sterile conditions to establish a GBM cell line through the following steps. First, we washed the biopsy with DPBS (Fisher Scientific, #14190250), then finely chopped it with a scalpel. Next, enzymatic dissociation was performed at 37°C for 10 minutes using TrypLETM enzyme (Thermofisher Scientific Gibco™, #12604013). To neutralize the enzyme, three volumes of DMEM (Thermofisher Scientific Gibco™, #11965118) supplemented with 5% KnockOut™ Serum Replacement (Thermofisher Scientific Gibco™, #A318502) was added. After that, cells were pelleted, and supernatant was trashed after a centrifugation at 1000 RPM for 4 minutes. GBM media was then added to the tube, and the pellet was dissociated with ten up-and-down pipettings, followed by another centrifugation at 300 RPM for 2 minutes. Both the supernatant and the pellet were placed separately into wells of a 6-well ultra-low adherence plate (Fisher Scientific, #07-200-602) with GBM media supplemented with 1x penicillin-streptomycin (Fisher Scientific, #15140148), incubating them at 37°C with 5% CO2. After a few weeks, neurospheres formed in suspension, contingent upon the presence of glioblastoma stem cells (GSCs) in the GBM biopsy. Finally, neurospheres were dissociated and cultured as monolayers in GBM medium without antibiotics, using wells coated with GeltrexTM (Gibco, #A14133).

### Limited dilution assay

On the day of the experiment, cells were seeded in an ultra-low attachment 96-well plate (Fisher Scientific Corning™ Costar™, #07-200-603) filled with *Neurosphere* media. The plate was divided into four sections, each containing 24 wells with 3, 10, 30, or 100 cells. After seven days, the number of negative wells were counted under a microscope. A negative well was defined as a well with no neurosphere formation. The experiment was performed in duplicate. By plotting the percentage of wells without neurospheres against the log of the number of cells deposited per well (3, 10, 30, 100), we generated a linear regression (y = ax + b). When plating at limiting dilutions, the proportion of culture wells with no neurospheres (negative wells) follows the zero term of the Poisson distribution: F0 = e-x. The number of cells needed to isolate one stem cell (x = 1) is calculated using F0 = e(−1) = 0.37 or 37%. The percentage of GSC cells was assessed using the following formula: % GSCs = (a/(ln(37) - b)) * 100.

### Metabolomic assay

For metabolomics, cells were processed using a standardized protocol [42,43]. In brief, after being passaged, each primary cell line was seeded in two 10-cm dishes. Two days later, samples of about 1,000,000 cells were harvested for metabolomics. To this end, cells were rinsed twice with ice-cold saline (NaCl 0.9%) on ice. After the second wash, cells were harvested in dry ice-cold 80% methanol and stored at −80°C. On the day of analysis, samples were extracted by sonication in ice-cold water before being derivatized by methoxymation and silylation. Samples were then analyzed using an Agilent 8890 GC with a DB5-MS + DG capillary column coupled to an Agilent 5977B MS using electron impact ionization (EI) at 70 eV (Agilent Technologies, Santa Clara, CA, USA). Finally, the Agilent MassHunter Workstation software was used for analysis and specific metabolites were identified using the NIST/EPA/NIH mass spectra library (NIST, Gaithersburg, MD, USA).

### RT-qPCR

Total RNA from cell pellets was collected using QIAshredder columns (QIAGEN, #79656) and the EZ-10 kit (Bio-basic, #BS88583). 2500 ng of RNA was treated with DNase (Thermo Fisher Scientific, #EN0521), and then 500 ng of treated RNA was reverse transcribed into cDNA using the iScript gDNA clear cDNA synthesis kit (Bio-Rad, #1725035). cDNA was quantified using the fluorescent probe SYBR-green (Applied Biosystems, #4472908), and qPCR was performed in triplicate. A pool of non-reverse transcribed RNA from each line served as a control sample. For RT-qPCR analysis, the relative expression of each gene was calculated in GBM lines using the delta Cq method. RPL27 was used as the housekeeping gene. A list of ARN targets is available in **Table 1**. Target genes were chosen according to a study published by Neftel et al. Identifying expression signatures correlated to different subtypes of cells present in GBM [5].

**Table 1.**
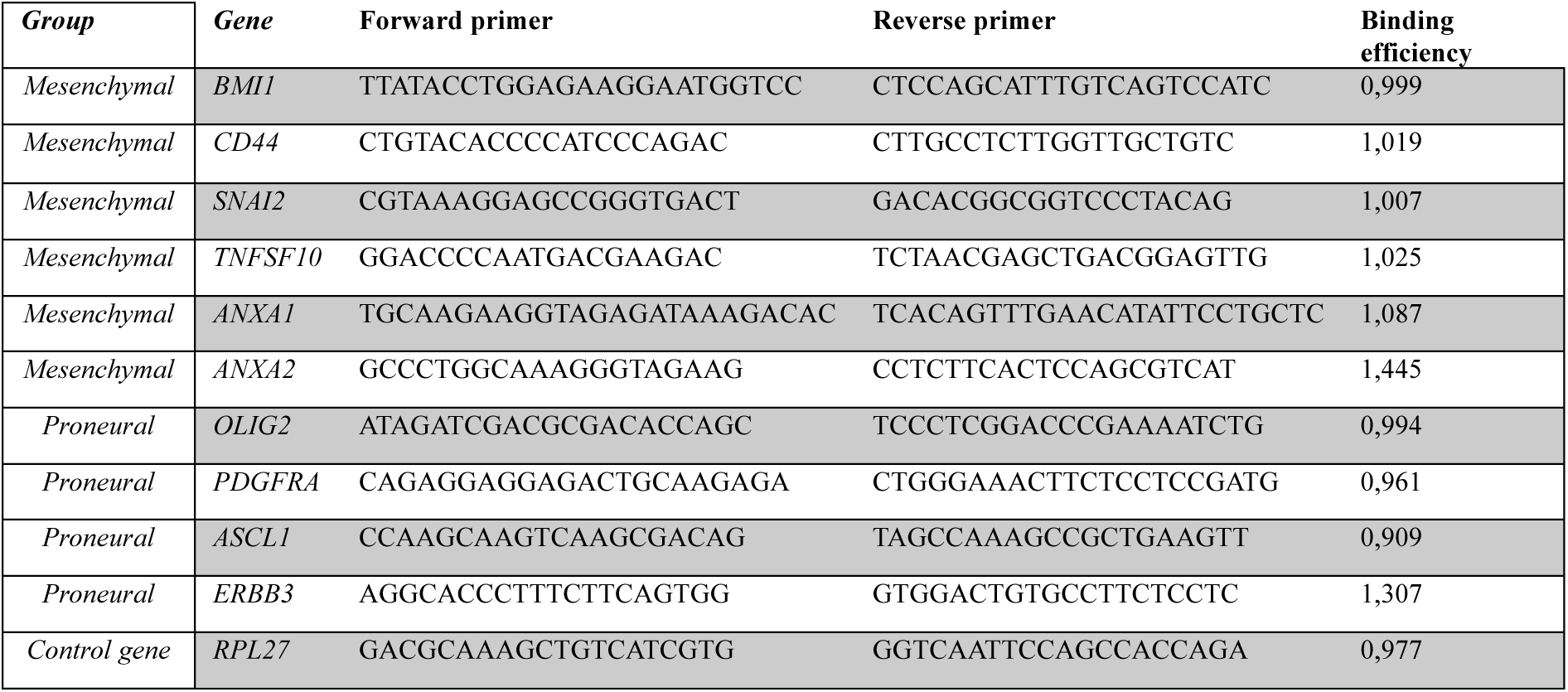
Characteristics of qPCR primer pairs.

### Statistical analyses

The D’Agostino & Pearson test was used to determine normality. When comparing two groups, Unpaired t-test was performed. In case data did not reached normality, a Mann-Whitney test was used. In figures representing pairwise comparisons, data are expressed as means ± standard error of the mean (SEM). All statistical analyses above were performed using GraphPad Prism 10.1.2 (GraphPad Software Inc., San Diego, CA, USA) or JMP Pro 17 (JMP Statistical Discovery LLC, Cary, NC, USA). Results were considered statistically significant when the p-value was less than 0.05.

### Principal component analysis

Dimension reduction was performed using principal component analysis (PCA). SRPlot was used to run the built-in ‘prcomp’ for PCA calculation and ‘FactoMineR R package’ for plotting the center and scale by default. The PCA biplot was calculated using ‘ggbiplot’ via SRPlot. Confidence ellipses (95%) were drawn around the groups to show the clustering patterns between the subtypes. Cluster heatmap was plotted using ‘pheatmap R package’ through SRPlot. Normalized abundance value was calculated using complete linkage clustering and the Euclidean distance method. The correlation coefficient plot was analyzed using standard statistical algorithms for Pearson’s correlation, and heatmaps were generated using SRPlot (https://bioinformatics.com.cn/srplot) [44].

## ACKNOWLEDGMENTS

The authors thank all the students and staff who assisted with sample collections at CHU de Québec – Université Laval, Hôpital de l’Enfant-Jésus, and especially the patients and their families for generously giving their time to participate in this study. S.M.I.H. and M.R. are supported by the Cancer Research Society (ID#1057382 to MR and SMIH; ID#1278359 to MEH). S.M.I.H. is a recipient of Junior 2 Research Scholar of the FRQS https://doi.org/10.69777/310750. E.A.W. holds the Canada Research Chair Tier 2 in Metabolic Vulnerabilities of Cancer and is supported by the Canada Foundation for Innovation (38622 and 45901) and the Cancer Research Society and the Suzanne and Louis Daubois fund (#1051060 and #1449872). S.M.I.H., M.R., E.A.W., and M.E.H. are supported by the Fondation du CHU de Québec. A.S is supported by Mitacs Elevate Postdoctoral Fellowship Program (#IT45505). This work was also supported by the Fonds de recherche du Québec (FRQ) through the Research Centre Grant. V.F. is a recipient of a training award of the Fonds de recherche du Québec-Santé (FRQS) https://doi.org/10.69777/291836. G.K. is a recipient of training awards from the Natural Sciences and Engineering Research Council of Canada (NSERC) and the FRQS https://doi.org/10.69777/307170. V.W. is a recipient of a training award of the Fonds de recherche du Québec - Nature et Technologies (FRQNT) https://doi.org/10.69777/271653.

## AUTHOR CONTRIBUTIONS

HLD: Conceptualization, Investigation, Formal Analysis, Methodology, Validation, Project administration, Writing—Review & Editing.

JM: Conceptualization, Investigation, Formal Analysis, Methodology, Validation, Writing— Review & Editing.

AS: Conceptualization, Methodology, Validation.

VF: Conceptualization, Methodology, Validation.

GK: Conceptualization, Methodology, Validation.

VW: Conceptualization, Methodology, Validation.

ML: Conceptualization, Methodology, Validation.

LB: Conceptualization, Methodology, Validation.

FR: Conceptualization, Methodology.

EB: Conceptualization, Methodology.

KM: Resources (recruitment of participants).

MD’A: Resources (recruitment of participants).

SS: Investigation.

MEH: Conceptualization, Validation, Project administration, Supervision

EAW: Investigation, Conceptualization, Formal Analysis, Conceptualization, Validation, Project administration, Writing—Review & Editing.

MR: Investigation, Conceptualization, Formal Analysis, Validation, Project administration, Funding acquisition, Supervision, Writing—original draft.

SMIH: Investigation, Conceptualization, Formal Analysis, Conceptualization, Validation, Project administration, Supervision, Writing—Review & Editing.

