## supplemental figures for "Glioblastoma gene expression based subtypes have defined metabolomic states"

### SUPPLEMENTARY MATERIAL

#### Supplementary Figure S1

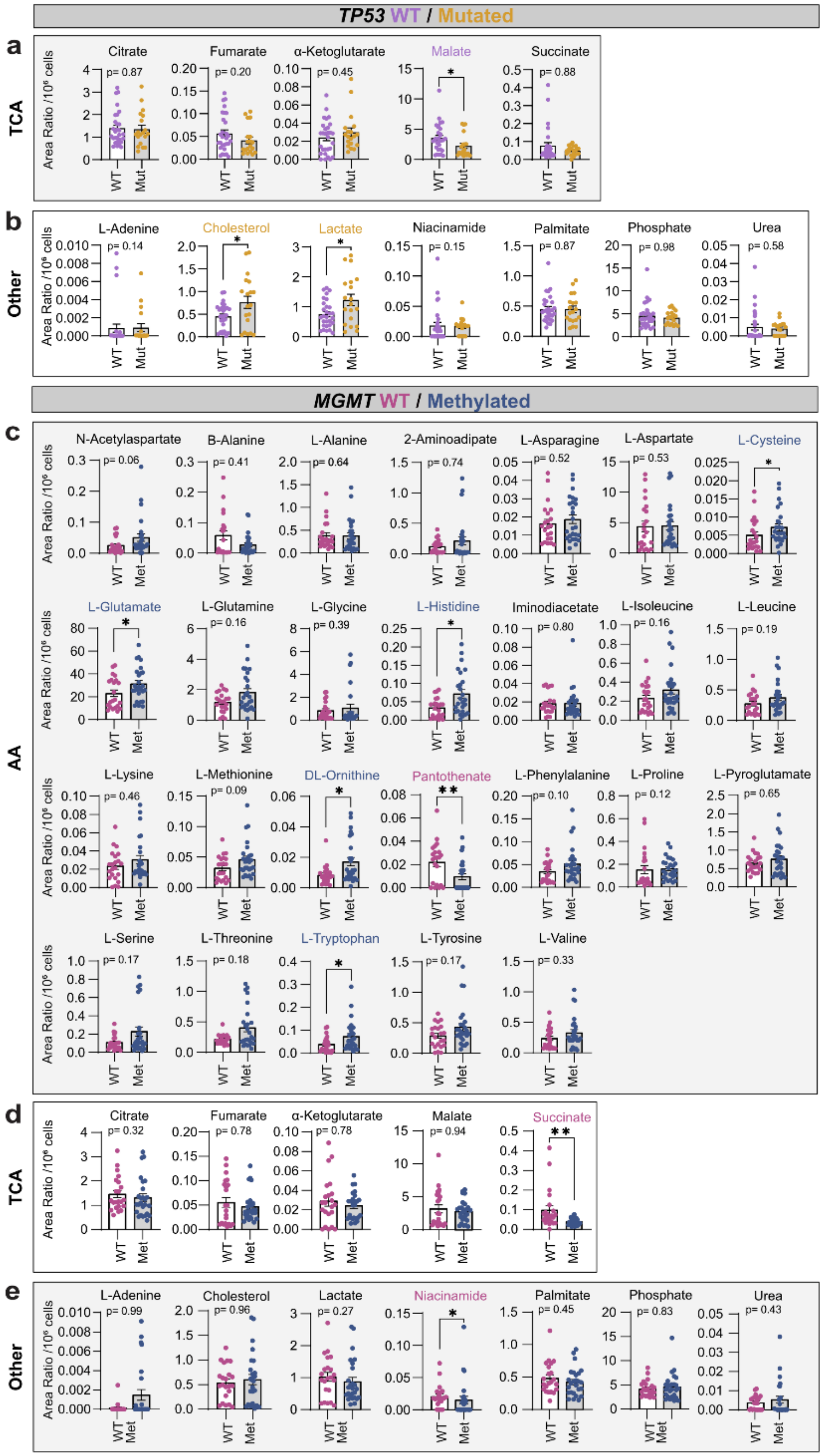

**Supplementary Figure S1. *TP53* mutations alters metabolomic profiles in GSC lines.** Comparison of TCA intermediates (**a**) and other metabolite (**b**) levels in cell lines with or without *TP53* mutations. Comparison of amino acids levels (**c**) as well as TCA intermediates (**d**) and other metabolite (**e**) in *MGMT* WT and *MGMT* met cell lines. *Statistical analyses:* Unpaired t-test or Mann-Whitney test. *Abbreviations:* AA, amino acids; Met, methylated; *MGMT*, $O^6$-methylguanine-DNA methyltransferase; Mut, mutated; TCA, Tricarboxylic Acid Cycle; *TP53*, tumor protein 53; WT, wild-type.

#### Supplementary Figure S2

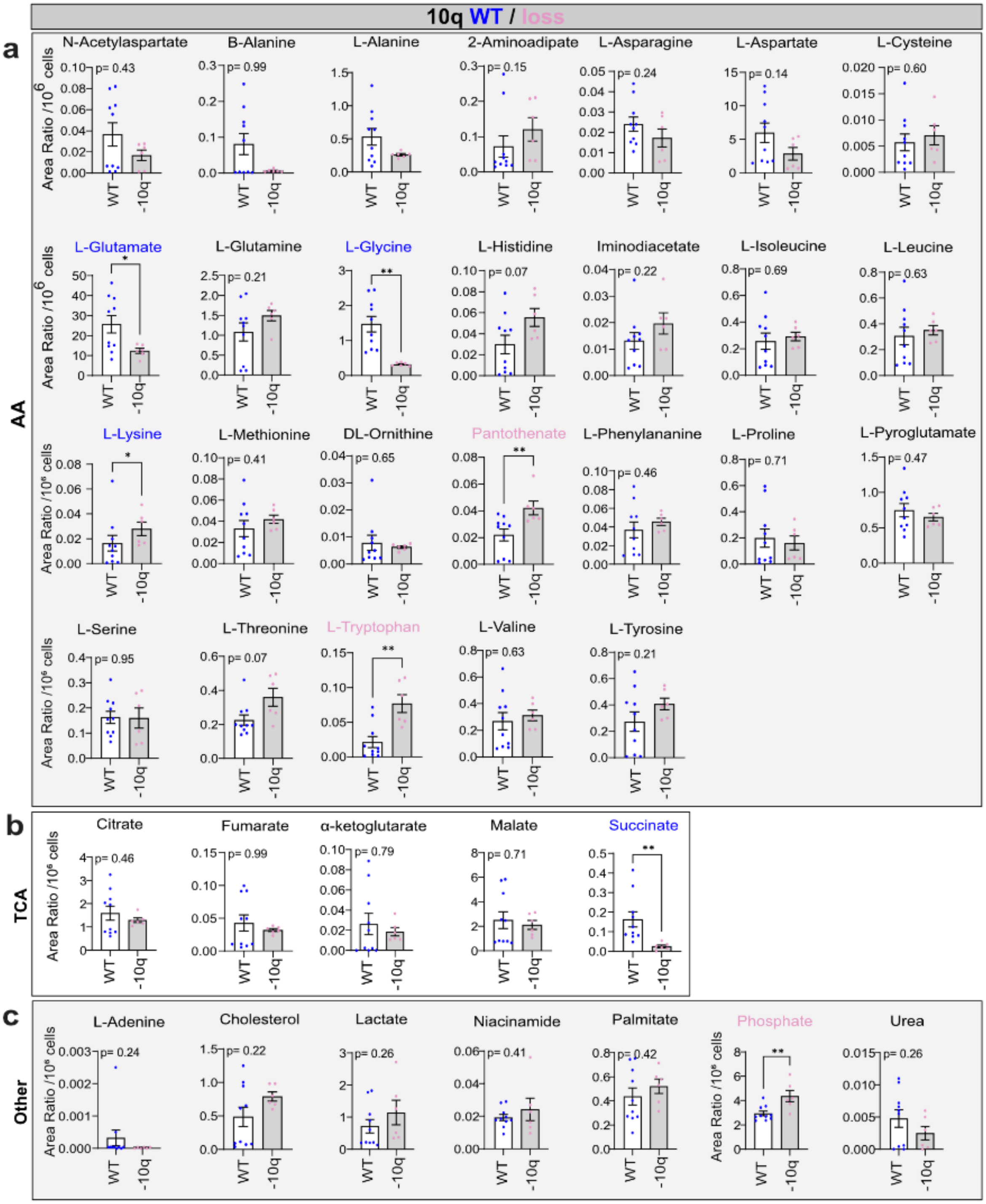

**Supplementary Figure S2. 10q loss status did not alter metabolomic profiles in GSCs lines.** Comparison of amino acids levels (**a**) as well as TCA intermediates (**b**) and other metabolite (**c**) in cell lines with or without 10q loss. *Statistical analyses:* Unpaired t-test or Mann-Whitney test. *Abbreviations:* AA, amino acids; TCA, Tricarboxylic Acid Cycle; WT, wild-type; -10q, loss of the chromosome 10q.

### Supplementary Figure S3

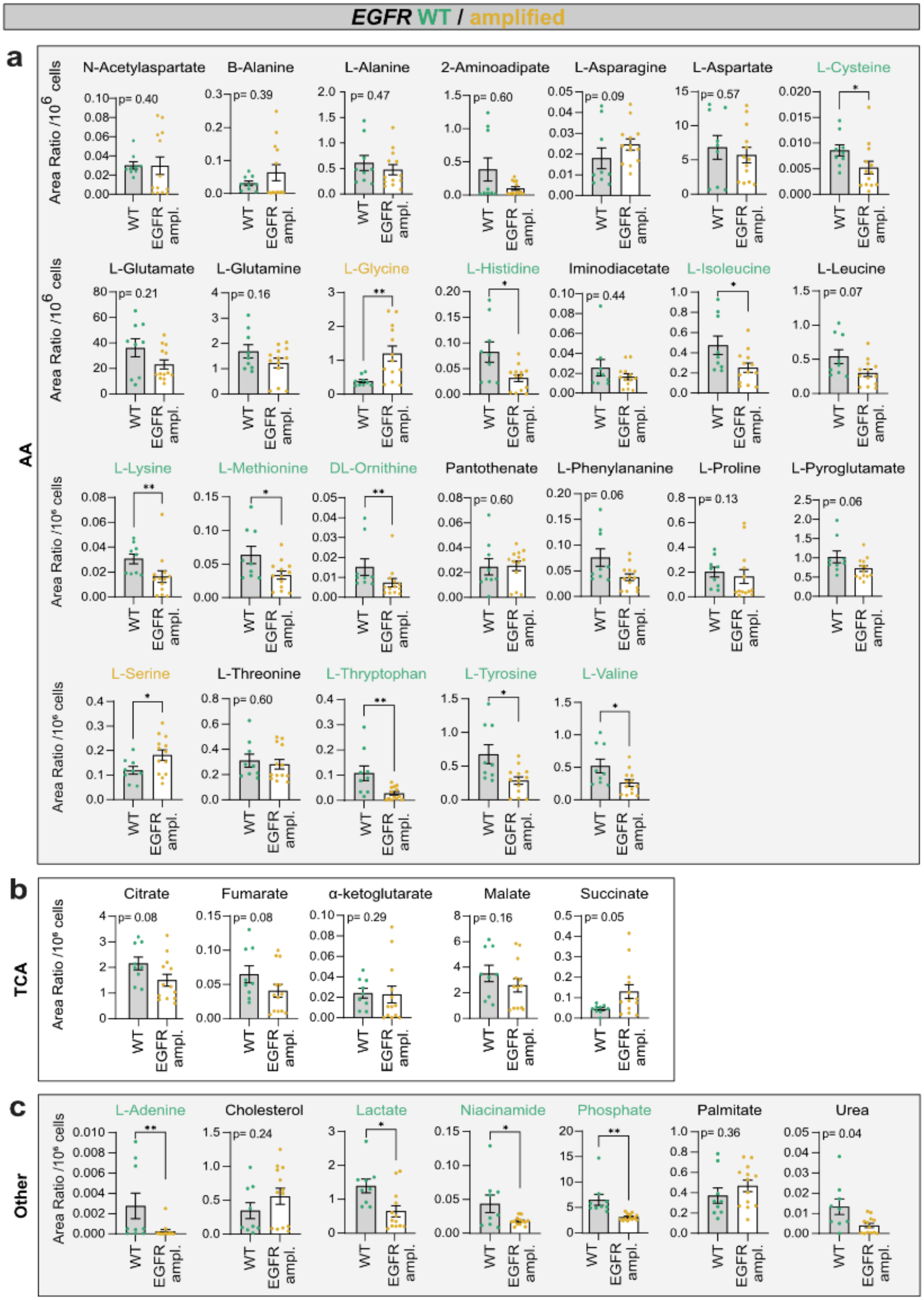

**Supplementary Figure S3.** ***EGFR*** **amplification status did not alter metabolomic profiles in GSCs lines.** Comparison of amino acids (**a**) levels as well as TCA intermediates (**b**) and other metabolite (**c**) in cell lines with or without EGFR amplification. *Statistical analyses:* Unpaired t-test or Mann-Whitney test. *Abbreviations:* ampl., amplification; AA, amino acids; *EGFR*, epidermal growth factor receptor; TCA, Tricarboxylic Acid Cycle; WT, wild-type.

#### Supplementary Figure S4

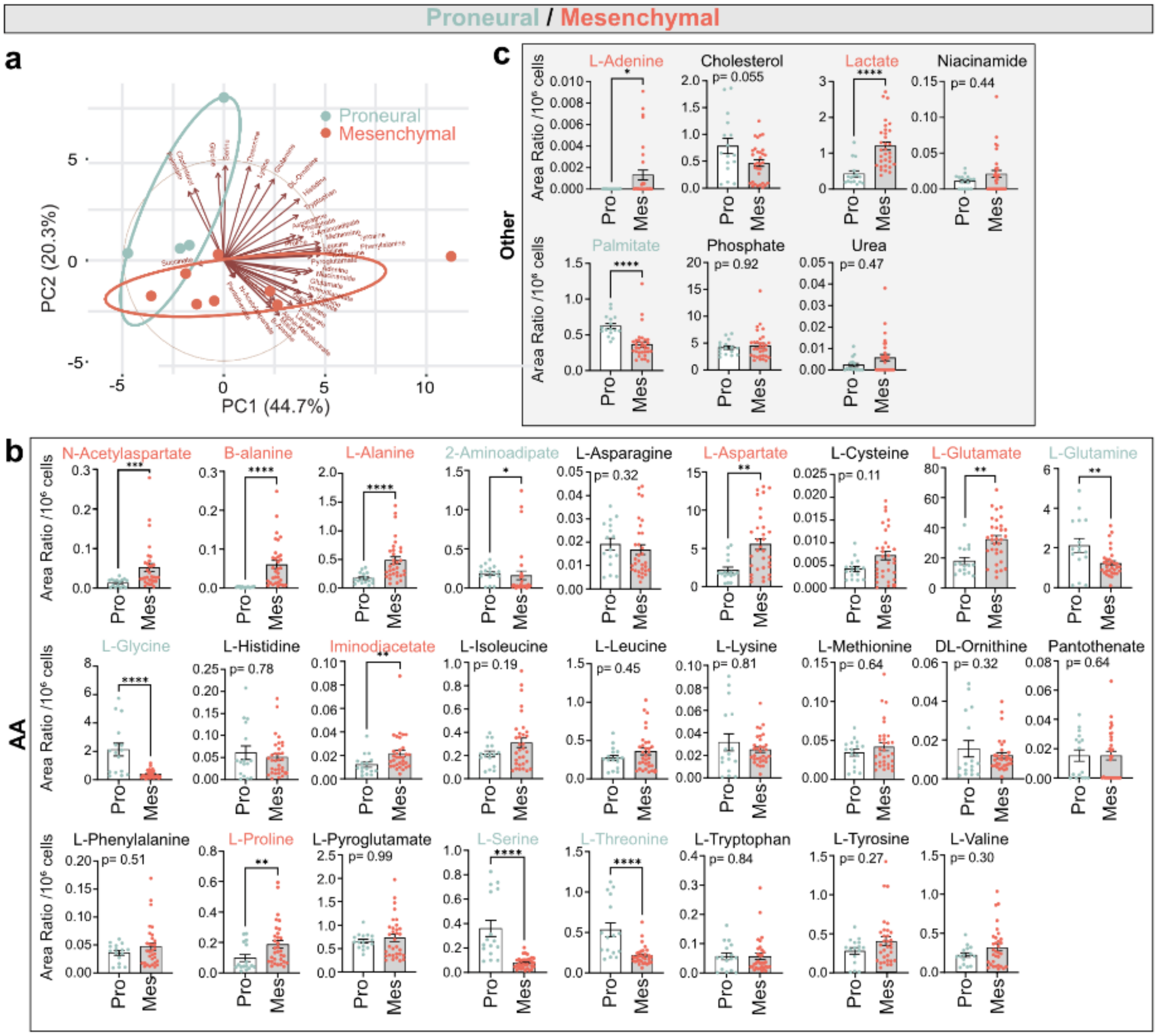


**Supplementary Figure S4. GBM subtypes show distinct metabolomic states. a.** PCA biplot showed that some metabolites levels are associated with proneural-like or mesenchymal-like cell lines. Comparison of (**c**) amino acids levels as well as other metabolite (**b**) levels in proneural-like or mesenchymal-like cell lines. *Statistical analysis:* **a.** Confidence ellipse (95%) highlights group clustering, and the arrows indicate the contribution and direction of individual metabolites to the observed separation between the subtypes. **b,c.** Unpaired t-test or Mann-Whitney test. *Abbreviations:* AA, amino acids; Mes, mesenchymal-like cells; Pro, proneural-like cells.
